# Palindrome Presence, Location, and Structural Context Are Predictive of RNA Self-Association

**DOI:** 10.64898/2026.11.05.764733

**Authors:** Changhwan Ahn, Haebin Park, Vailankanni L. Rodrigues, Gregory L. Dignon

## Abstract

RNA self-association is a key elementary event behind many cellular processes from ribosome assembly, to regulation of gene expression, to assembly of biomolecular condensates. While RNAs self-interact in many ways, one direct and unambiguous sequence-encoded pattern for RNA self-recognition is a palindrome, or palindromic motif. In isolation palindromic motifs tend to fold back on themselves, favoring stable stem-loop secondary structures. However, palindromes have been implicated in some cases of RNA self-recognition, segregation and phase separation, raising the question of which palindromes in a given RNA are actually available for intermolecular engagement. Toward answering this question, we define three classes of palindromic regions based on local secondary-structure context, and degree of availability for interactions via their nucleobases, sorting into three classes. We then apply this framework across known structures of RNA chains and find that the majority of palindromic regions are bound up in secondary structures. We then designed a minimal set of short stem-loop RNA sequences to evaluate the self-interaction stability of each defined palindrome class using coarse-grained molecular dynamics simulations. Overall, this work provides a residue-level, structure-aware framework that connects RNA sequence, secondary-structure context, and intermolecular self-association for two interacting RNA chains.

## Introduction

RNA is a structurally and functionally versatile biopolymer. Beyond its classical role in the central dogma as the intermediate carrier of genetic information [11, 10], RNA serves as a translational adapter (tRNA), a majority component of the ribosome (rRNA), and a regulator of gene expression through diverse non-coding species [7, 16, 20, 8]. Underlying this functional breadth is a chemistry that supports both intramolecular folding and intermolecular pairing. RNAs are linear polymers of ribonucleotides, and are capable of forming three-dimensional structures, similar to proteins, however, their limited nucleotide alphabet, and lack of true hydrophobic moieties limits their structural, and catalytic capacity[18]. Nevertheless a significant fraction of structures in the protein databank contain RNA molecules bound to proteins, multi-RNA complexes, or even single RNA molecules[4, 19]. Toward forming structures, each nucleotide can engage in Watson–Crick (A–U, G–C) or wobble (G–U) base pairs, and the resulting hydrogen-bonding patterns allow RNA to fold into hairpins, internal loops, and higher-order architectures, or to hybridize with a separate RNA strand [24, 32].

When two RNA molecules interact in trans, the resulting interactions underlie a wide range of cellular processes. Sequence-specific RNA–RNA recognition mediates the silencing of mRNAs by microRNAs and small interfering RNAs, the modification of ribosomal and other RNAs by snoR-NAs, and the regulation of transcript stability and translation by long non-coding RNAs [3, 29, 17]. At larger length scales, intermolecular RNA–RNA interactions serve as multivalent scaffolds that contribute to the assembly of biomolecular condensates and membraneless organelles, such as P-bodies, stress granules, and nucleoli, which organize biochemistry without a lipid boundary [5, 28, 33, 34]. The material properties of these condensates are influenced by the sequence-encoded architecture of the RNA molecules they contain [35, 26]. While RNA self-association spans a vast spectrum from individual microRNA–mRNA pairs to multi-component condensates, these sequence-encoded interactions ultimately dictate the biophysical properties of the local cellular environment. To better understand the event mechanistically, we ask how we focus on this elementary case and ask how the primary sequence and the local secondary-structure context of an RNA chain together encode its self-specificity, that is, its tendency to selectively pair with a second copy of itself rather than with an arbitrary partner.

To evaluate this event mechanistically, we investigated whether an RNA sequence favors intramolecular self-association over intermolecular pairing. Although RNA self-association is well-established, we focus our attention on a unique feature of these interactions and investigate palin-dromic regions. A palindromic sequence is inherently reverse-complementary, allowing an RNA chain to pair either with its own sequence or with a neighboring strand.

Among the sequence motifs that can specify such an interaction, palindromic regions are the most direct. A palindromic region is one whose reverse complement matches itself, so that two copies of a palindrome meet the Watson–Crick complementarity requirement for homotypic duplex formation without any need for an external template, providing a sequence-encoded self-recognition motif for two RNA chains. Palindromes have long been recognized as biologically significant in DNA, where they form the recognition sites of type II restriction enzymes and contribute to recombination, replication, and the sequence-dependent shape of the double helix [6, 23]. However, the mere presence of a palindrome is insufficient to predict its intermolecular engagement. In a single-stranded RNA context, an isolated palindromic region carries a strong thermodynamic tendency to fold back on itself, maximizing stable intramolecular base pairs [24, 32]. Consequently, the same sequence feature that endows RNA with trans self-specificity also makes it prone to cis sequestration — a balance that, at the level of an individual RNA molecule, contributes to whether a given palindromic region is available for intermolecular pairing.

In a given RNA, then, a palindrome is not simply “present” or “absent”; it occupies a specific structural context that contributes to whether its constituent bases are thermodynamically accessible to a second RNA chain. To formalize this, we classify each palindromic residue, on the basis of its local secondary-structure environment, into one of three distinct classes. A palindromic residue belongs to **class 1 (Hairpin)** if it is contained within a stem-loop structure, having base pairs between residues within a single palindrom. It belongs to **class 2 (Comb)** if it is itself engaged in an intramolecular base pair, but not with other residues within the same palindromic region. These base pairs can be on one side of a local hairpin stem or between long-range pairs far apart in the RNA. The remaining unpaired residues, those not enclosed by an apical loop (e.g., residues in internal bulges, asymmetric loops, or free termini), are assigned to **class 3 (Strand)**.

Building on this classification, we examine RNA palindromes at three complementary levels. We first apply a linear-time algorithm to enumerate palindromic regions across the human transcriptome (GENCODE [13]) and experimentally resolved RNA structures from the Protein Data Bank, using x3dna-DSSR [22] for secondary-structure annotation, and characterize their length distribution and per-residue structural-class composition. We then probe the thermodynamic consequences of structural context by performing coarse-grained temperature replica-exchange molecular dynamics simulations [2, 26, 31] on a small set of short RNA sequences designed to place a single extended palindrome into each of the three structural classes, summarizing the resulting monomer and dimer melting behavior using multiple linear regression models built on the structural-class residue counts. Finally, we use two-dimensional free-energy surfaces to map intramolecular folding against intermolecular binding at the residue level. Together, these analyses provide a residue-resolved framework that links primary RNA sequence, secondary-structure context, and intermolecular self-association for two interacting RNA chains in our coarse-grained model system.

## Materials and Methods

### Datasets and preprocessing

To compare the sequence-level prevalence of palindromic motifs against their distribution in experimentally resolved structures, we curated two datasets, one having comprehensive sequence information from the RNA transcriptome, and one having sequence and structural information from the Protein Data Bank(PDB). The transcriptomic dataset was obtained from GENCODE release 49 (v49) [13, 25] and comprises approximately 5.34 × 10^5^ human transcripts spanning all annotated biotypes (protein-coding, lncRNA, retained intron, NMD, pseudogene, and others). The structural dataset was assembled from RNA chains in the Protein Data Bank (PDB) [4], screened for high resolution X-ray crystal structures with reported resolution ≤ 3.5 °A, or solution NMR structures. We also filtered out RNA sequences shorter than 20 nucleotides, yielding a final dataset comprising 8,062 individual RNA molecules.

The structural dataset was further split into subsets based on the composition of the parent PDB entry: RNA–protein complexes (*N* = 6,440), RNA–RNA complexes (*N* = 1,137), and single-RNA structures (*N* = 485). Sequence-level analyses (palindrome detection, length distributions, and density statistics) were performed on the full GENCODE and PDB datasets; secondary-structure-based analyses, including the per-residue structural classification, were performed only on the PDB dataset, for which experimentally resolved structures provide reliable secondary-structure assignments.

### Secondary-structure assignment

For experimentally resolved RNA chains in the PDB, secondary-structure annotations in dot-bracket notation were derived directly from the 3D atomic coordinates using the x3dna-DSSR software suite [22]. DSSR identifies base pairs from physical hydrogen-bonding patterns and base-stacking geometry; for the present analysis we considered the canonical Watson–Crick A–U and G–C pairs together with the wobble G–U pair. Other detected interactions were not included in our analysis for this study. Secondary structure of RNA strands was processed as dot-bracket strings, highlighting location and partnering of residues in base pairs[14]. Visualization of secondary structure was done using forna, through the ViennaRNA package[15, 21].

### Palindrome detection algorithm

To identify self-complementary regions across large datasets, we implemented a roughly linear-time (∼ *O*(*n*)) palindrome detection algorithm. The algorithm scans the sequence for nucleation dinucleotides, defined as any contiguous dinucleotide that forms a valid base pair (A–U, U–A, G–C, C–G, G–U, or U–G). Upon detecting a valid dinucleotide at positions *i* and *i* + 1, the algorithm symmetrically extends the alignment outward to positions *i*−*k* and *i*+1+*k* for increasing *k*, stopping when complementarity is broken or the sequence boundary is reached. For the length-distribution analysis, nested palindromic subsequences were recorded (i.e. a 6-nt palindrome contains a 4-nt and a 2-nt self-complementary subsequence), for comparison to random expectation. For the density and per-residue classification analyses, only maximal palindromes were retained.

#### Palindrome statistics in transcriptome and theoretical baseline

Under our definition of a palindrome, the probability that a random RNA sequence of even length *N* is a palindrome can be calculated as:

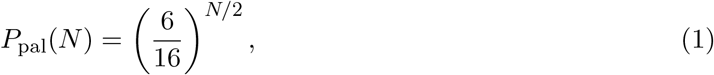

where the factor 6*/*16 accounts for the six valid base-pairs out of sixteen possible dinucleotide sequences. The exponent *N/*2 arises because only the first half of a palindromic sequence is independent, while the second half is determined by the complementarity constraint.

We apply this statistical framework to full RNA sequences by enumerating the expected prevalence of palindromic regions within all possible windows of the sequence. The expected number of *n*-residue palindromes in a transcript of length *L* is

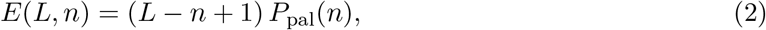

where we multiply the probability of a random *n*-nucleotide window being a palindrome by the number of length-*n* windows (*L* − *n* + 1) in the sequence. For a 1,000-nucleotide RNA, this predicts approximately *E*(1000, 10) ≈ 7.35 palindromes of length 10.

For per-residue analyses we use two complementary metrics. The *palindrome count per nucleotide* (palin/nt) is the total number of maximal palindromes in a sequence divided by its length *L*. The random-baseline expectation for palin/nt is equivalent to the expected number of 2-residue palindromes, because any longer palindromes will always have shorter palindromes nested inside. Thus our random expectation for number of maximal palindromes in a sequence of length L is,

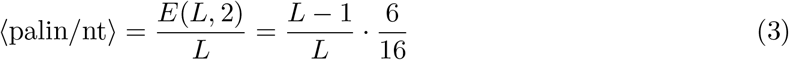

This converges to 0.375 at relatively short chain lengths (*L* ≳ 20).

The *palindromic residues per nucleotide* (palin res/nt) is the cumulative number of nucleotides participating in any maximal palindrome divided by *L*. Summing across all even palindrome lengths and weighting each by the *n* residues it contributes gives the random-baseline expectation

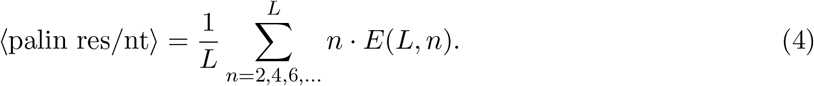

For sufficiently long random sequences, Eq. 4 yields a baseline expectation of approximately 1.20 palindromic residues per nucleotide. The value exceeds unity because a single residue can participate in multiple palindromic regions at the same position. Equations 1–4 provide the analytical reference against which we compare the palindrome statistics measured on biological RNAs.

### Coarse-grained molecular dynamics simulations

#### Force field and model parameters

We used a single-bead-per-nucleotide coarse-grained representation in which each nucleotide is modeled as a single interaction site at the phosphate position with mass 330 Da, adapted from Aierken *et al*. [2, 26]. The model includes harmonic bonded and angle energy terms for sequential nucleotides in a chain. It also includes excluded-volume interactions to prevent nonphysical chain overlap, and a sequence-specific base-pair interactions (*U*_base pair_) encoding Watson–Crick and wobble pairing geometry. Simulations were run using the LAMMPS simulation software [31].

To mitigate kinetic trapping in the rugged folding-and-binding landscape, we used temperature replica-exchange molecular dynamics (T-REMD) [30]. Sixteen replicas were geometrically distributed across the temperature range 150.00–600.00 K, spanning the folding–unfolding transition of all six designed sequences. Simulations were performed in LAMMPS [31] in the NVT ensemble using Langevin dynamics (*T*_damp_ = 100 ps) and with replica exchanges between adjacent temperatures attempted every 10,000 steps. Each replica was integrated for 2.5 × 10^8^ steps at a 10 fs timestep (2.5 *μ*s per replica) and the first 10 ns of each replica trajectory were discarded as equilibration. Single-chain simulations used a cubic simulation box of side length 200 °A; two-chain simulations used a cubic box of side length 250 °A, with the two chains initialized in random, non-overlapping orientations. To estimate statistical uncertainty, we used block averaging by dividing each trajectory into 5 contiguous blocks of equal length, and the standard error of the mean was computed as 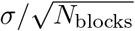, where σ is the standard deviation of the 5 block means.

#### Melting temperature determination

Melting temperatures *T*_*m*_ were obtained by fitting the temperature-dependent fraction of intact native base pairs, *f* (*T* ), to a four-parameter sigmoidal function:

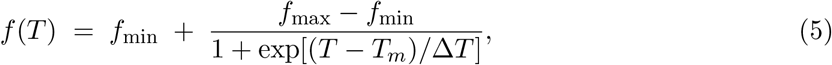

where *f*_min_ and *f*_max_ are the lower (unfolded) and upper (folded) baselines, *T*_*m*_ is the inflectionpoint melting temperature, and Δ*T* characterizes the width of the transition. Fits were performed with non-linear least squares.

#### Free-energy surface analysis

For each sequence and each replica temperature, we computed the joint probability distribution *P* (*N*_intra_, *N*_inter_) from the equilibrated portion of the trajectory by histogramming. The free-energy surface was obtained from the inverse Boltzmann relation,

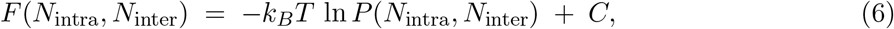

where *k*_*B*_ is the Boltzmann constant, *T* is the simulation temperature, and *C* is chosen so that *F*_min_ = 0 for each panel. Free-energy values are reported in units of *k*_*B*_*T*.

#### Statistical analysis and visualization

Data analysis and visualization were performed in Python using `numpy, scipy, pandas, seaborn, and matplotlib`.

## Results

### A minimal palindromic RNA dimerization system

RNA palindromes are nucleotide sequences characterized by self-complementarity, in which the sequence read 5^′^ to 3^′^ on one strand is complementary to itself in reverse. Unlike linguistic palindromes (e.g., “racecar”), which rely on character symmetry, RNA palindromes are defined by complementarity, rather than identity. In this work, we consider palindromes that are self-complementary following the standard Watson–Crick A–U and G–C pairs as well as the wobble G–U pair.

We begin by considering a minimalist system of a single short RNA sequence that is completely palindromic (Figure 2). When two copies of this sequence come into contact, they can fully hybridize, forming a dsRNA helix (Fig. 2, right). When unbound, the two copies of the palindromic RNA will generally be most stable as a stem-loop “hairpin” structure with a loop of four unpaired residues.The free energy of binding is influenced primarily by the number of base pairs formed and by the translational and rotational entropy of the two chains. The hybridized dsRNA duplex is energetically favored, having four additional base pairs relative to the sum of the two hairpin stems, but entropically disfavored, since both chains lose significant translational and rotational entropy (Fig. 2). We do not anticipate a large difference in configurational entropy, since both cases involve relatively stable ordered structures. The hybridization of a purely palindromic RNA sequence can therefore be tuned by temperature, which influences the relative importance of entropy versus energy, and by RNA concentration, which influences the amount of translational entropy lost upon binding.

**Figure 1:**
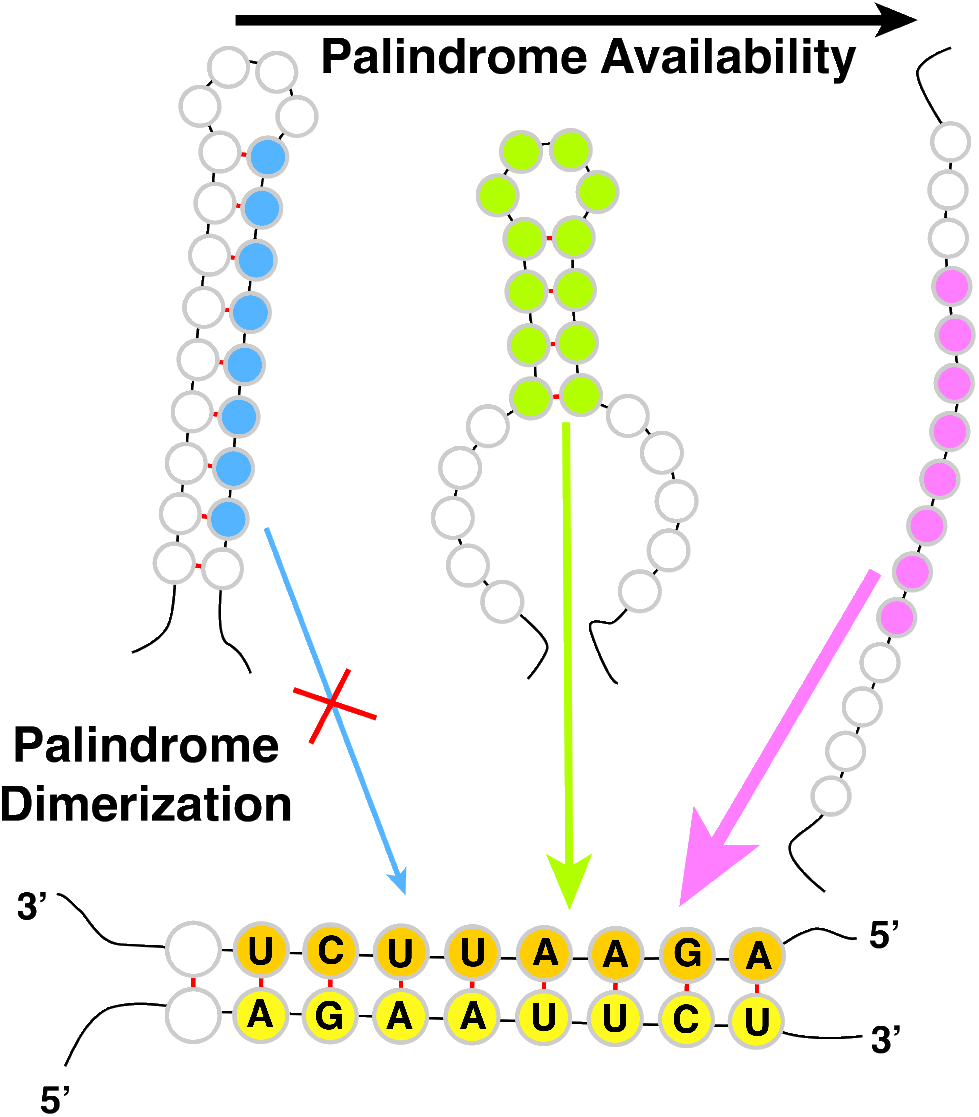
TOC Figure

**Figure 2:**
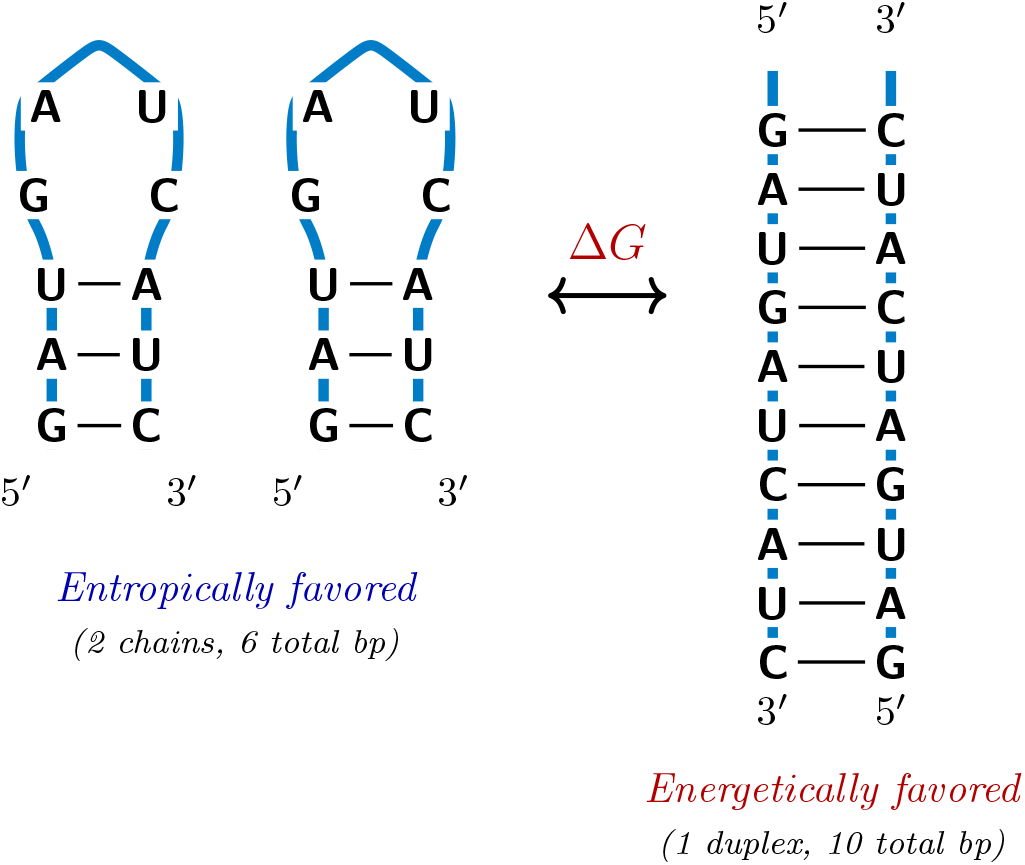
Schematic of palindromic RNA dimerization. A representative dimerization equilibrium for a fully palindromic 10-nt RNA sequence (5^′^-GAUGAUCAUC-3^′^). *Left:* two separate copies of the same chain, each folded into a stem-loop hairpin and *Right:* the same two chains hybridized in antiparallel orientation into a fully paired duplex.

### Transcriptome-wide prevalence of palindromes

Having defined RNA palindromes, we next ask whether palindromes are present in known RNA sequences. Theoretically, long palindromic sequences should be rare, exponentially less likely at longer lengths (Eq 1). Using a comprehensive transcriptomic dataset of over half a million human RNA sequences, we detect the number and length of all palindromes within each sequence, and compare the prevalence as a function of palindrome length (Fig. 3). For comparison, we also analyzed a significantly smaller dataset of RNA sequences present in the protein databank. In each palindrome length value, the prevalence is normalized by the total number of possible windows of that length in the full list of sequences.

**Figure 3:**
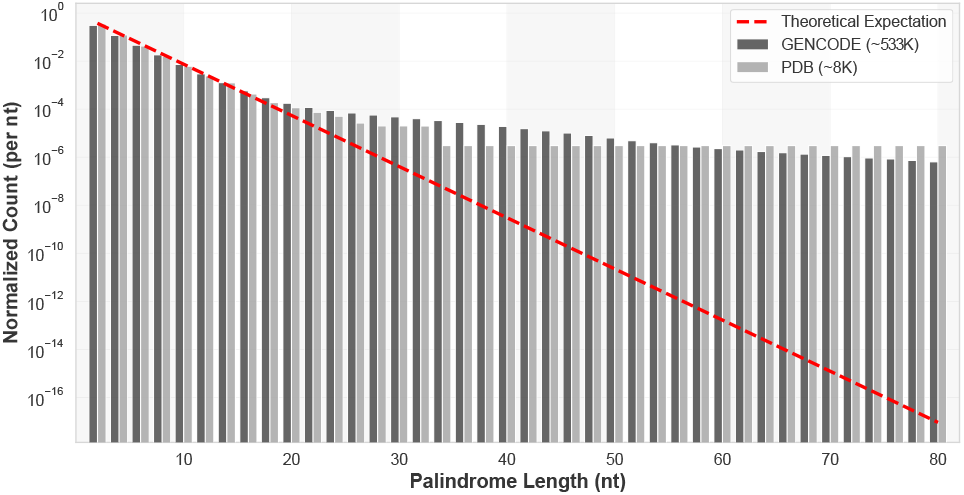
Normalized palindrome counts per nucleotide are plotted against palindrome length for the GENCODE human transcriptome (black), PDB structures (grey), and the theoretical random expectation (red, Eq. 1).

The random expectation (red) follows a strict exponential decay that appears linear on a semi-logarithmic scale, providing a baseline in which short palindromes (≲ 10 nt) are statistically common while longer palindromes are vanishingly rare (Fig. 3). Both biological datasets follow this baseline at short palindrome lengths but deviate from it as length increases. The human transcriptome (black) exhibits a heavy tail for palindromes longer than ∼20 nt, with frequencies orders of magnitude above the random expectation. Since both show enrichment of longer palin-dromes, it seems plausible that there is some selective pressure for long palindromes in nucleotide sequences[27, 9]. It is important to specify that though the prevalence for longer palindromes is sometimes many orders of magnitude higher than the random expectation, it represents only a relatively small number of very long palindromes. This pressure could be functionally related to RNA self-association, but may also reflect an indirect contribution from DNA-level palindromic motifs such as transcription factor binding sites or other regulatory elements [12], which could be passively transcribed into RNA. Regardless of their origin or function, the presence of these long self-complementary regions would carry the consequence that any RNA containing them would have some sequence-encoded potential for self-association and self-specificity.

Beyond the question of longer palindrome prevalence throughout the full datasets, we also considered the fluctuations in numbers from individual RNA strands. We define the metric palin-drome count per nucleotide (palin/nt, eq. 3), which tracks the total number of maximal palindromes within a sequence, ignoring nested palindromes within a longer palindrome. A second complementary metric is the number of palindromic residues per nucleotide (palindromic res/nt; eq. 4), which measures the cumulative fraction of nucleotides that participate in palindromic motifs. The first gives a rough approximation of how many independent sites in an RNA strand could potentially form self-specific intermolecular interactions, and the second also factors in the average size of these self-specific regions.

We applied these metrics to our two RNA sequence datasets, and found that in both cases, we get relatively wide distributions, centered close to the random expectation (Fig. 4A,B). Firstly, it is clear that the GENCODE dataset is a much narrower distribution than the PDB dataset, which is likely due to the presence of longer sequences, which would lead to smaller deviations from expectation within individual chains (i.e. the law of large numbers). The average length of sequence in GENCODE is 1151 nucleotides, while the average sequence from PDB is only 58. Additionally, both metrics fall slightly below the theoretical random expectation, indicating that while many sequences have significant palindromic content, there seems to be some selective pressure against their presence in RNA transcripts. Importantly, this does not conflict with the previous result that long palindromes are significantly enriched beyond random expectation because the total number of these long palindromes is still very small compared to the total number of shorter (i.e. ≤ 20 nt) palindromes. We conclude therefore that of the known RNA transcripts, individual transcripts have a wide variety of palindromic content, from significantly less to significantly more than random expectation. It is plausible to speculate that higher or lower palindromic content could be useful functionally. This has also validated that the sequences selected from the protein databank have overall similar palindromic content to the overall RNA transcriptome.

**Figure 4:**
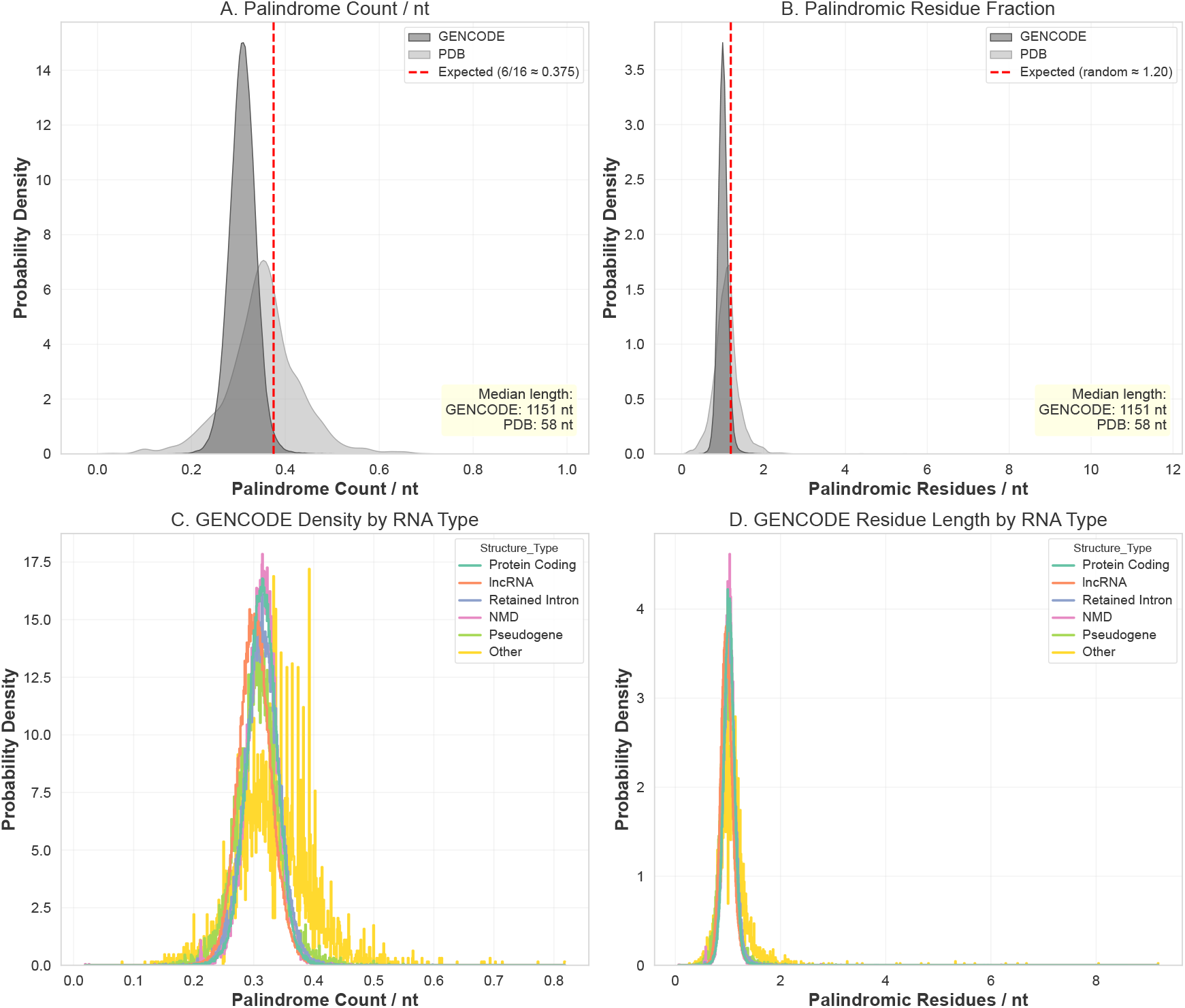
Palindrome statistics in the GENCODE transcriptome and PDB structural dataset. (A) Palindrome count per nucleotide (palin/nt) and (B) palindromic residues per nucleotide (palin res/nt) for the full PDB (*N* = 8,062,) and full GENCODE (*N* = 533,704) datasets. (C–D) The same metrics broken down by GENCODE biotype.

**G**To further investigate the prevalence of palindromes in different RNAs, we decomposed the GENCODE dataset to subsets by the included classifying biotypes, including protein-coding mR-NAs, long non-coding RNAs (lncRNAs), retained introns, NMD transcripts, and pseudogenes.Each class seems to follow a similar distribution to each other, with minimal deviation except in the “Other” category. Taken together, the two density metrics suggest that palindromic content, at the level of overall density, is a generic feature of cellular RNA sequences rather than a property confined to any single functional class; the values sit modestly below the random-sequence baseline in both the structural and transcriptomic datasets. Further studies may look more deeply into palindromic content vs. different constitutively, or functionally-related RNA sequences, but that is beyond the scope of this work.

### Structural classification: the availability hypothesis

Detecting a palindrome confirms its potential for self-complementarity but does not, on its own, indicate whether the palindrome is readily available for intermolecular binding. We must further consider the position within an RNA structure, or structural ensemble. Towards this, we define three distinct classes for a given palindromic residue, based on its local secondary-structure context. Figure 5 illustrates the three classes on a model RNA sequence secondary structure.

**Figure 5:**
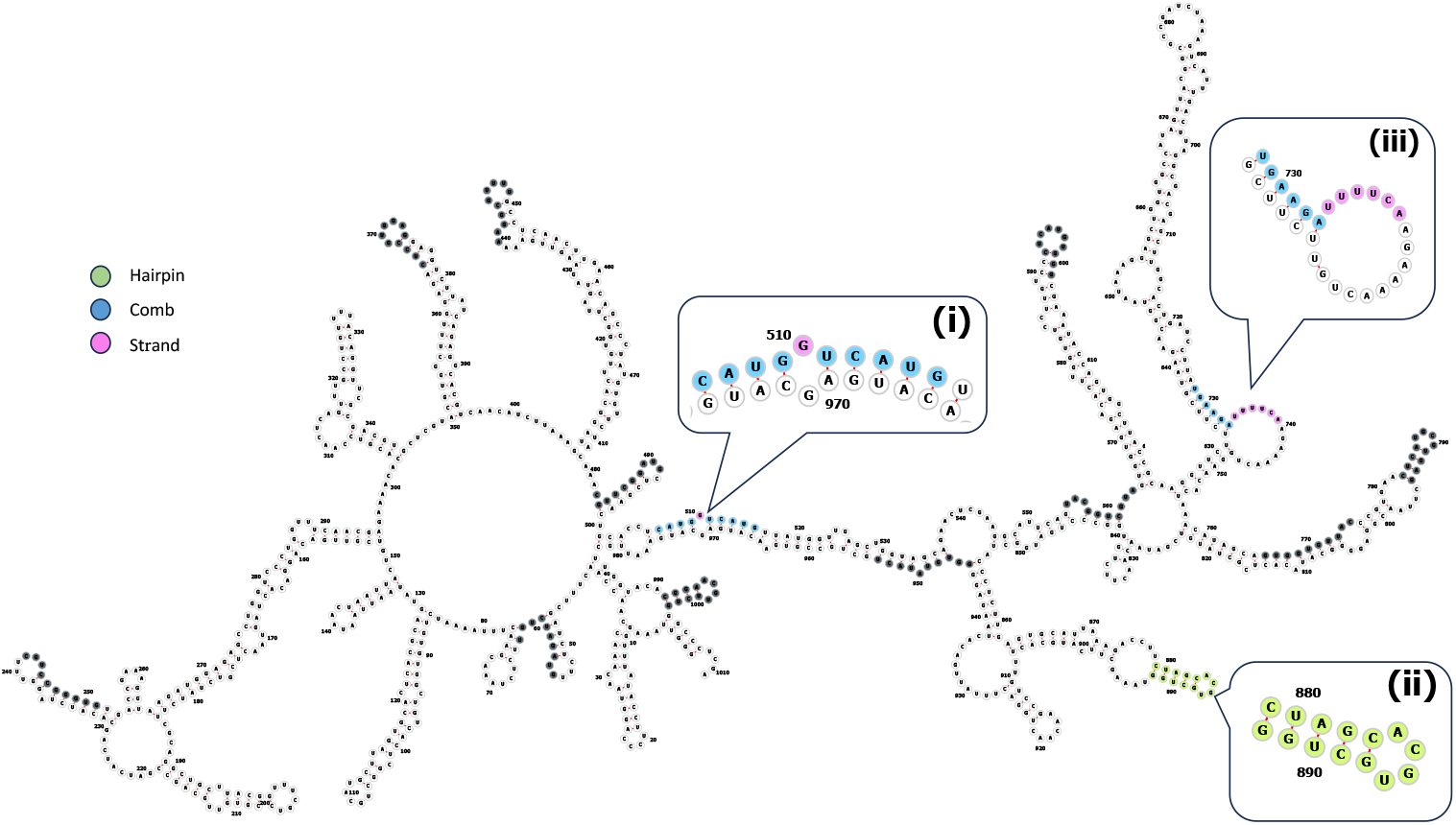
Structural classification of palindromic residues within an example structured RNA architecture (predicted secondary structure of the first 1,000 nucleotides of the SARS-CoV-2 genome). Palindromic residues are highlighted in gray, and insets show examples of different classes of palin-dromes.

**Class 1 (Hairpin)** comprises palindromic residues that fully reside within a stem-loop structure of a structured RNA (Figure 5, green). In this configuration, both arms of the stem are engaged in intramolecular base pairs and are therefore locally “closed,” while the residues within the loop itself are unpaired and somewhat available for cross interactions with other bases. Thus, class 1 palindromes have a mixed character, where many of the bases are bound up to other bases within the palindrome, and having four bases at the loop being partially accessible. Importantly, this is the most stable structure for a palindromic sequence in isolation (Fig. 2), but is not the only possible state for palindromic regions in longer sequences that aren’t fully palindromic (Fig. 5).

**Class 2 (Comb)** comprises palindromic residues that are actively engaged in intramolecular base pairing, but not with other bases contained in the same palindrome (Figure 5, blue). Notably, class 2 includes residues forming both local hairpin stems and long-range structural scaffolds, but always involve only one side of a stem unlike class 1. These also have the capacity to be more structurally stable than class 1 palindromes since they can form stable base pairs at every base, without needing an unpaired loop region.

**Class 3 (Strand)** comprises palindromic residues residing in unpaired regions that are not part of a stem, or the loop in a stem-loop structure. This includes internal bulges, asymmetric loops, long loops, and free termini (Figure 5, red). Lacking base-pairing partners, class 3 residues have the lowest barrier to participate in intermolecular base pairing, having no need to disrupt local secondary structure. We note, however, that fully extended class 3 palindromes are uncommon in practice since self-complementary sequences of more than ∼8 residues have a strong intrinsic tendency to fold back on themselves into hairpins.Thus, it seems most palindromes are a combination of classes, which is why our following analysis focuses on palindromic residues of each class, rather than full palindromes.

We then turn to the pdb dataset to analyze the palindrome prevalence, and structural classification of palindromes in known RNA structures. We first analyze all structures, and find that, in general, palindromic residues in the second class are the most common, while those in hairpin configurations are the least common (Fig. 6A). We further split the dataset into three subsets, including RNA-protein complexes, multi-RNA complexes, and single RNAs, and find that the overall number of palindromes is consistent between each, but the number of palindromic residues, i.e. length of palindromes differs somewhat between the subsets (Fig. S1). Within these subsets, we continue to find that the class 2 palindromic residues are most common, followed by class 3 and then class 1 (Fig. 6B-D).

**Figure 6:**
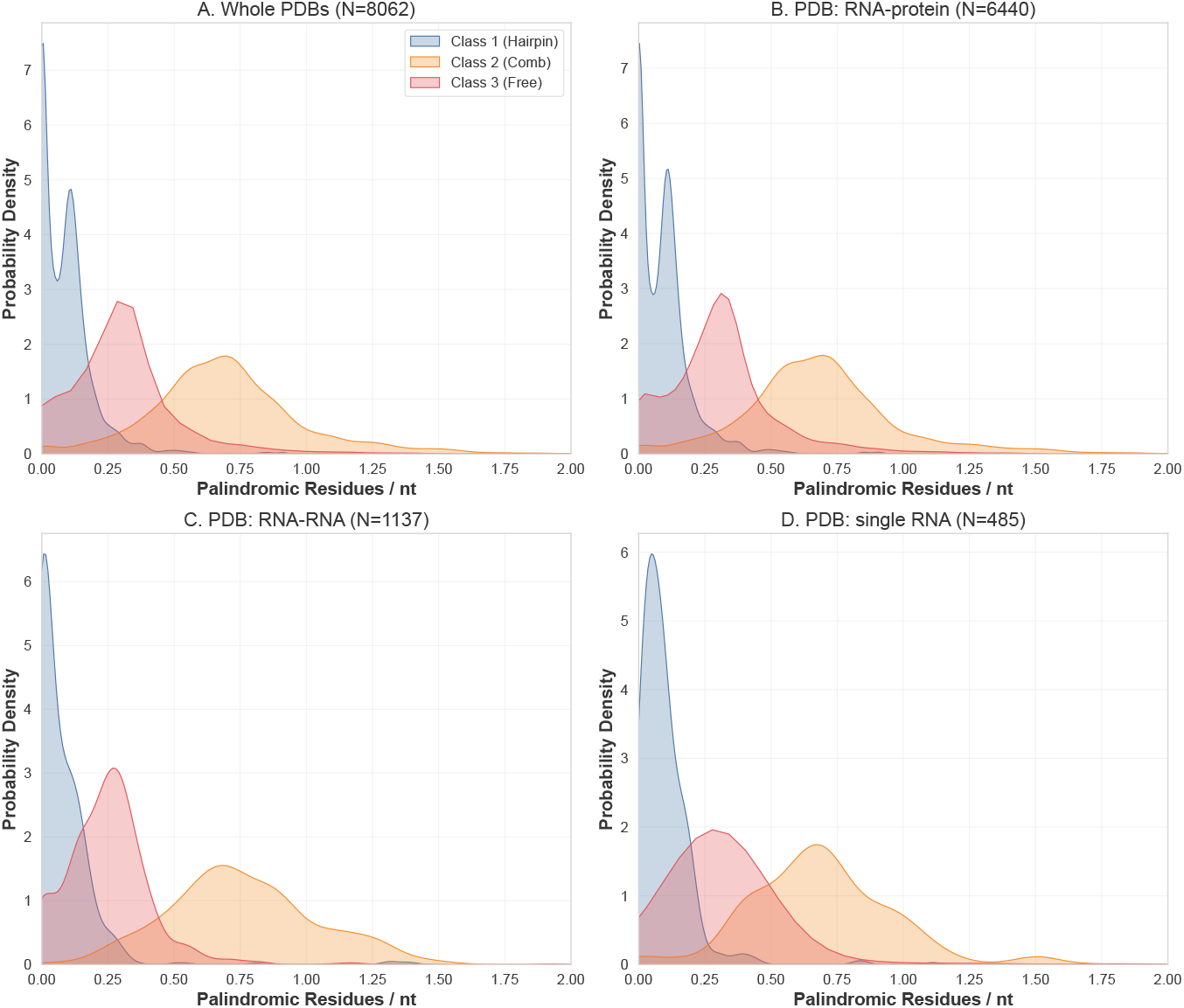
Distributions of the three palindrome structural classes within experimentally resolved RNA structures from the PDB. Probability densities of palindromic residues per nucleotide for (A) the entire PDB dataset, (B) RNA–protein complexes, (C) RNA–RNA complexes, and (D) single-chain RNA structures.

It is hard to draw conclusions about selective pressure for any one class due to the different functions served, and challenges with normalizing to many different RNA sequences. We interpret this as showing that, of the self-complementary regions in RNA sequences, the majority of them are tightly bound up into class 2, comb-like structures, and are less available for interactions with other copies of themselves. However, there is still a significant fraction of “free” class 3 palin-dromic residues in many sequences, which indicate that many RNAs could have self-specificity at such regions. Finally, we note that this is merely a statistical observation about the distribution of palindrome contexts in the PDB, and does not, on its own, demonstrate that class 3 palin-dromic residues drive intermolecular association in any specific RNA. To examine the underlying thermodynamics directly, we turn to coarse-grained molecular dynamics simulations.

### Thermodynamic analysis of the availability hypothesis

#### Design of model sequences

We next investigate the thermodynamics of the competition between intrachain folding, and interchain binding of palindromic regions in RNA. Toward this, we designed six Single-Palindrome Hairpin sequences, termed SPH sequences, as minimal systems to represent the different classes of palindromes. These were designed to contain only a single extended (≥ 6 nt) palindrome placed different structural contexts according to our previously-defined classes (Figure 7). Another important feature is that these sequences were designed to minimize the prevalence of alternate, “non-native” configurations. We validated the prevalence of the designed secondary structures using AlphaFold2, RNAFold, and MD simulations, as discussed below[1, 21].

**Figure 7:**
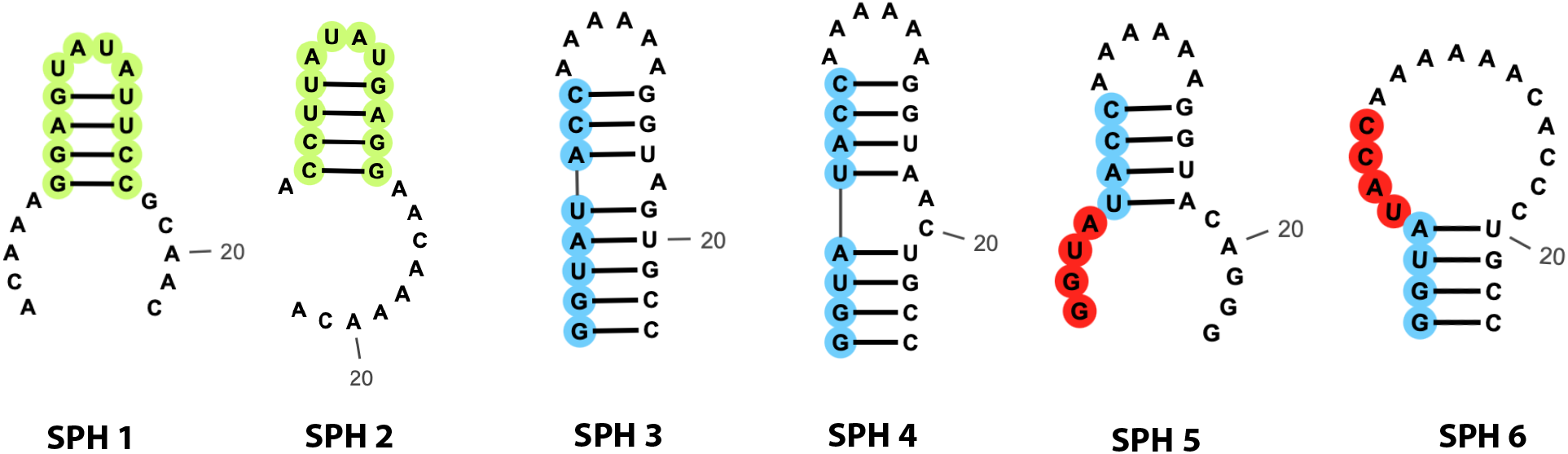
Six designed RNA sequences spanning the three structural classes. SPH1–2 each include self-contained stem-loops, SPH3–4 contains a palindrome as part of a stem, and SPH5–6 include a palindrome that is partially located in an unpaired loop region.

The first two sequences, SPH1-2, each contain a class 1 palindrome, which is achieved by embedding the palindromic region within a self-contained stem-loop which folds back on itself. These two sequences differ from each other only by the location of the hairpin in the sequence. For SPH1, the palindrome is centered in the sequence, while SPH2 has it toward the 5’ terminal, which we expect to have a greater entropic loss upon folding. The next two sequences, SPH3-4, place the palindromic region on one side of a double-stranded stem formed by base-pairing with a complementary region elsewhere on the same strand, making them class 2 palindromes. Notably, both sequences contain mismatches on the complementary strand, in the form of a single, or two-residue bulge in SPH3 and SPH4 respectively. To have a fully complementary second strand would, by necessity, introduce a second palindrome into the sequence, and thus, within our design constraints, it would be impossible to design an SPH sequence with only a single perfectly base-paired class 2 palindrome. Finally, the last two sequences, SPH5-6 were designed to have a partial class 3 palindrome. Having a fully class 3 palindrome over 8 nt long is not possible since any fully unpaired palindromic region will begin to fold in on itself like a (class 1) hairpin. In both SPH5 and SPH6, we have an 8 residue palindromic region where half of its bases are paired (class 2) and half are unpaired (class 3). We note that the SPH6 sequence has a much longer loop than SPH5, and the paired bases are all toward the termini of the sequence. Both of these factors strongly oppose folding of the sequence, and thus, we will find that SPH6 represents the closest practical approximation to a fully exposed class 3 palindromic region within our design criteria.

Each of these sequences differ in quantifiable ways which may influence folding stability, two-chain self-interactions, or both. Toward building an understanding of the molecular determinants of folding and binding stability, we first extract a list of sequence and structure-based quantities for each of our designs. These parameters are tabulated in Table 1.

**Table 1:**
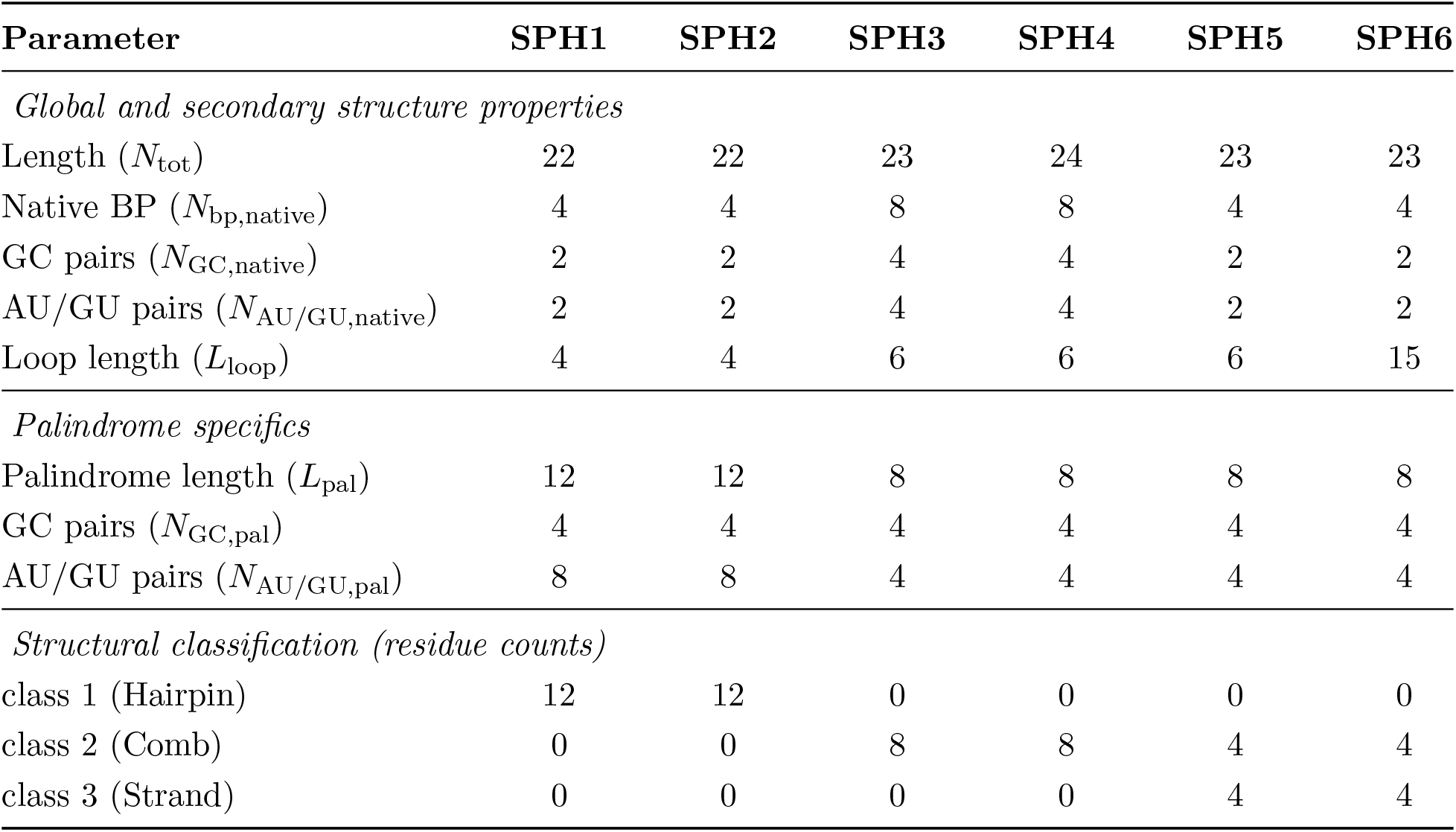
Physical and structural properties of the six designed RNA sequences.

#### Single-chain folding stability

We first assessed the intrinsic folding stability of each SPH sequence in isolation by conducting coarse-grained T-REMD simulations. This approach yields good sampling of temperature-dependent behavior and stability of the predicted secondary structure of the RNAs, which we quantify as the fraction of native intramolecular base pairs, *f*_intra_ (Figure 8A). The temperature range used is nonphysical for experimental studies, but is not expected to correspond precisely with real temperatures. Rather, our intention is to qualitatively compare the folding stability of our designed sequences.

**Figure 8:**
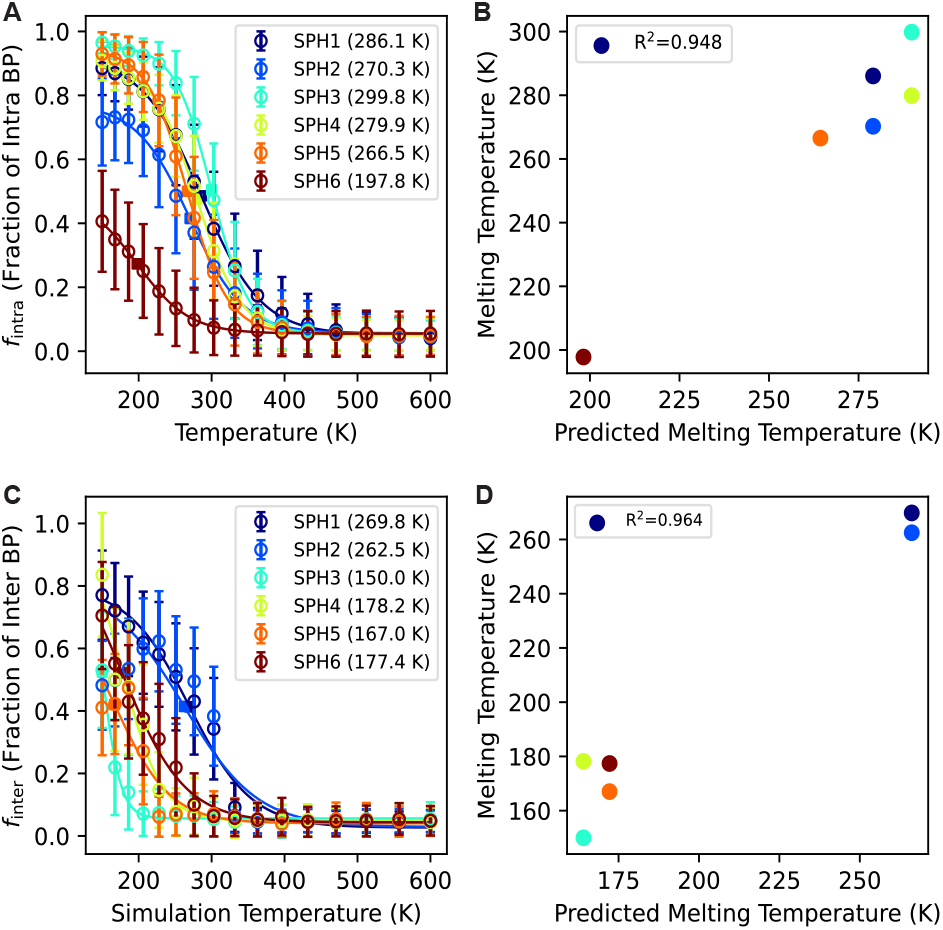
Single-chain folding and two-chain binding for SPH sequences from coarse-grained T-REMD simulations. A) Melting curves for single-molecule simulations. Scatter points are from simulation, lines are sigmoidal fit. B) MLR-predicted versus observed 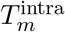 from Eq. 7. C) Fraction of intermolecular palindrome base pairs, *f*_inter_(*T* ). D) MLR-predicted versus observed *T*_*m*,*dimer*_from Eq. 8 (*R*^2^ = 0.964). Error bars represent the standard error of the mean from block averaging.

After calculating the fraction of native base pairs for each sequence, we fitted the melting curves to a sigmoidal function (eq. 5), and extracted the melting temperatures *T*_*m*,*fold*_ for each case (Fig. 8A,B). The first two sequences SPH1-2 yield relatively high melting temperatures, where SPH1 is somwhat more stable than SPH2, as expected from the effects of chain entropy loss close to the termini of the sequence. We find that the class 2 palindromic regions, SPH3-4 appear to have the greatest folding stability, where SPH3 is the highest melting temperature overall, and SPH4 is only slightly lower than that of the class 1 SPH1 sequence. Notably, the higher melting temperature for SPH3-4 may be due to the larger number of base pairs than SPH1-2, rather than the location of the palindrome. However, the presence of bulges and mismatches in the stem would then reduce the folding stability to offset the increased stability of more base pairs in the stem. This is clearly demonstrated by SPH3 which has only one mismatch and a higher melting temperature, compared to SPH4 which has two mismatched residues, and a lower melting temperature. SPH5 (class 3) yields a slightly lower melting temperature compared to the other sequences. Notably, we would not expect the location of the palindromic region to have a significant influence on the melting temperature of a single chain. This is observed here since the non-palindrome-related properties of SPH5 are nearly identical to SPH1, except that it has a loop region with six residues, rather than Finally, SPH6 carries a substantially longer unstructured loop of 15 residues and yields by far the lowest *T*_*m*,*fold*_ = 197.8 K, consistent with a large entropic penalty for loop closure outweighing the limited enthalpic gain from its short stem. This also highlights SPH6 as a sequence having a pure, class 3 palindromic region at most temperatures.

Using this dataset, we summarize the structural stability of our sequences by predicting *T*_*m*,*fold*_ through a multiple linear regression (MLR) model with minimal parameters:

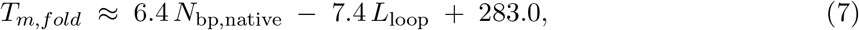

Our model fits the observed *T*_*m*_ values with *R*^2^ = 0.948 (Figure 8B) using only two parameters. The positive coefficient on *N*_bp,native_ is consistent with each native base pair contributing enthalpic stabilization, while the negative coefficient on *L*_loop_ (K per loop nucleotide) is consistent with the entropic cost of loop closure. We did not include additional parameters (e.g., the number or size of internal bulges) because the dataset comprises only six sequences, and adding further predictors would carry a high risk of overfitting. Other potentially-important parameters include the fraction of native base pairs that are G–C as they will have a greater stabilizing effect than other types of base pairs. Within our dataset, we cannot decouple this from *N*_bp,native_ because all SPH sequences have half of the base pairs as G–C.

#### Two-chain binding and palindrome structural context

To extend our investigation into competition of single-chain folding and intermolecular binding, we performed additional T-REMD simulations of two copies of an RNA strand. We quantified binding using the intermolecular base-pair fraction *f*_*inter*_(*T* ), defined as the number of cross-chain palin-dromic contacts normalized by the maximum possible intermolecular base pairs (Figure 8C). Fitting each curve to a sigmoidal two-state function yielded the dimer melting temperatures *T*_*m*,*dimer>*_.

Intriguingly, we found that SPH1 and SPH2 yielded the highest dimer melting temperatures by a significant margin (Fig. 8C,D). One significant contributor to this effect would be the longer palindromic regions (*L*_*pal*_ = 12) compared to the other four sequences (*L*_*pal*_ = 8). The most intriguing result is that SPH3, which had the highest *T*_*m*,*fold*_ has the lowest *T*_*m*,dimer_, overall, lower than we could estimate with confidence. This, along with the similarly low *T*_*m*,*dimer*_for SPH4 is consistent with our hypothesis that class 2, comb-shaped palindromic regions are the least available for cross-interactions with other molecules. We also consider the class 3 containing sequences, SPH5 and SPH6, finding that they generally have higher affinity for intermolecular interactions than the class 2 sequences. SPH6 is no longer the least stable of the two, and seems to prefer intermolecular binding over intrachain folding.

The two-chain interaction results are less intuitive than the single chain results. We thus fit an MLR model with a few parameters to see how each correlates with *T*_*m*,*dimer*_:

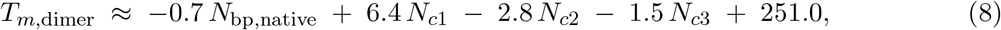

The agreement with simulated *T*_*m*,*dimer*_is quite good, with *R*^2^ = 0.964 (Figure 8D). We selected this particular set of four predictors deliberately to test whether the structural class composition of a sequence carries information about its dimer stability that is not already captured by *N*_bp,native_alone.

The signs of the regression coefficients are consistent with the picture in which existing intramolecular structure penalizes intermolecular binding. The negative coefficient on *N*_bp,native_is consistent with the same native base pairs that stabilize the monomer acting as a barrier to dimerization. The strongly negative coefficient on *N*_*c*2_ is consistent with class 2 (Comb) residues providing a particularly large barrier, since each such residue is engaged in an intramolecular base pair and would need to be released for the palindrome to bind in trans. The negative coefficient on *N*_*c*3_ is small and reflects the limited population of class 3 residues across this sequence set. The positive coefficient on *N*_*c*1_ is perhaps surprising, but may be confounded by the longer palindrome length of the class 1 sequences. We therefore do not interpret the *N*_*c*1_ coefficient as a generic property of class 1 architectures across all RNAs. Most importantly, we identified that the number of class 2 palindromic residues (*N*_*c*2_) seems to oppose intermolecular interaction more strongly than other factors, particularly more so than the total number of base pairs in the native structure, *N*_bp,native_.

### Competition of folding and binding

To more directly observe the competition and mechanism of folding and binding, we plot the melting curves for both single-chain and two-chain simulations, as well as the folding curves for each individual chain in the two-chain simulation (Fig. 9). Here we observe that for both class 1 hairpin sequences, SPH1-2, the dimer formation and fold are competing with each other (Fig. 9A,B). This is sensible because the same palindromic residues are forming base pairs in both the folded structure, and the hybridized two-chain dimer structure, and cannot participate in both. In both SPH1 and SPH2 at the conditions tested here, we find that inter-chain palindrome-palindrome hybridization is favored over intrachain hairpin formation, and this is demonstrated by the significant drop of native-like base pairs in the two-chain simulation (grey) compared to the single-chain simulation (black). Notably, the melting temperature of the dimer structure is dependent on concentration of the RNA, and thus introduces a possible design strategy to devlop RNA sequences that prefer folding or binding at particular temperatures, and can compete only in certain temperature regimes. The concentration used for the two-chain simulation in this case results in a dimer melting temperature comparable to the single-chain unfolding temperature.

**Figure 9:**
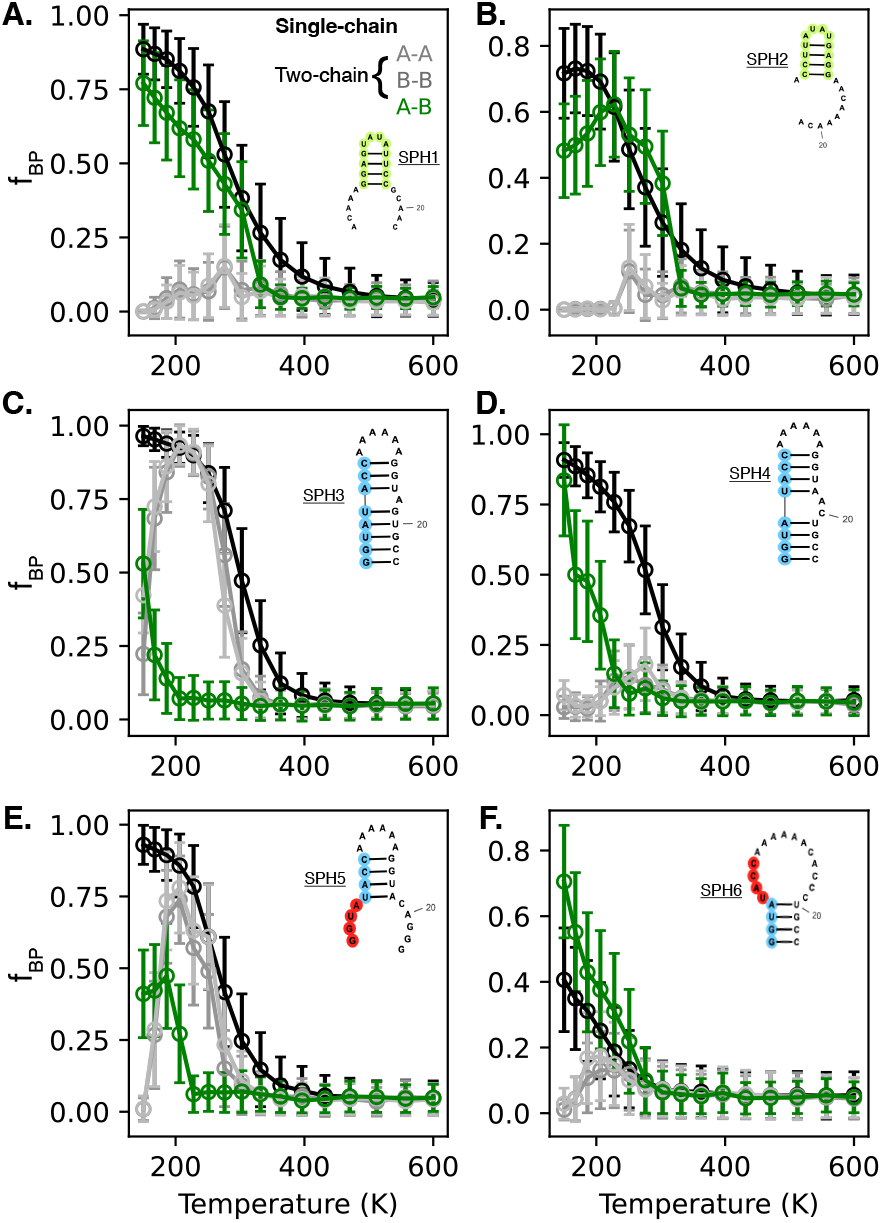
Competition between intramolecular folding and intermolecular binding for SPH1–6. Intramolecular base pairs from single-chain simulations are shown alongside intramolecular base pairs for each individual chain in the two-chain simulation (grey), and the intermolecular contact fraction (green). Note that the number of native base pairs is different than the number of interchain palindromic base pairs in each case, and fractions are normalized to the appropriate number.

A qualitatively different story arises when considering the class 2 palindromic sequences SPH3 and SPH4. Since these sequences were constructed to have greater native structure stability, and isolation of their palindromic regions, we anticipate the folded structure being stabilized and in-hibiting formation of the dimer structure. This is indeed observed in the data where the native-like folded structure of SPH3 is stabilized in both the single-chain and two-chain simulations and the two chains only form palindrome-palindrome intermolecular interactions at very low temperatures (Fig. 9C). The next case of SPH4 does not show this behavior however, and is likely the case because the native-like structure is disrupted by the 2-residue bulge on one side of the stem, which is significantly less stable than the 1-residue bulge in SPH3 (Fig. 9D).

The sequences with partially exposed class 3 palindromic regions show more unique behavior. SPH5 retains most of its folded structure when in 2-chain simulations, but at very low temperatures, palindrome-palindrome contacts begin to be partially favored, and seem to compete out folded native intra-chain base pairs (Fig. 9E). Fascinatingly, we find that the two-chain intermolecular binding is the strongest interaction in the case of SPH6, which has a very low folding fraction at all temperatures, and thus its palindromic regions are much more free to interact with other chains (Fig. 9F).

To further visualize the underlying mechanism, and possible conformations for each case we projected total contact fractions for intrachain, and inter-chain base pairs into two-dimensional free energy surfaces (Fig. 10; Fig. S3-8). We use a reference temperature of 251.1 K as it lies within the transitional regime of all six sequences in our coarse-grained model and therefore yields the most informative comparisons.

**Figure 10:**
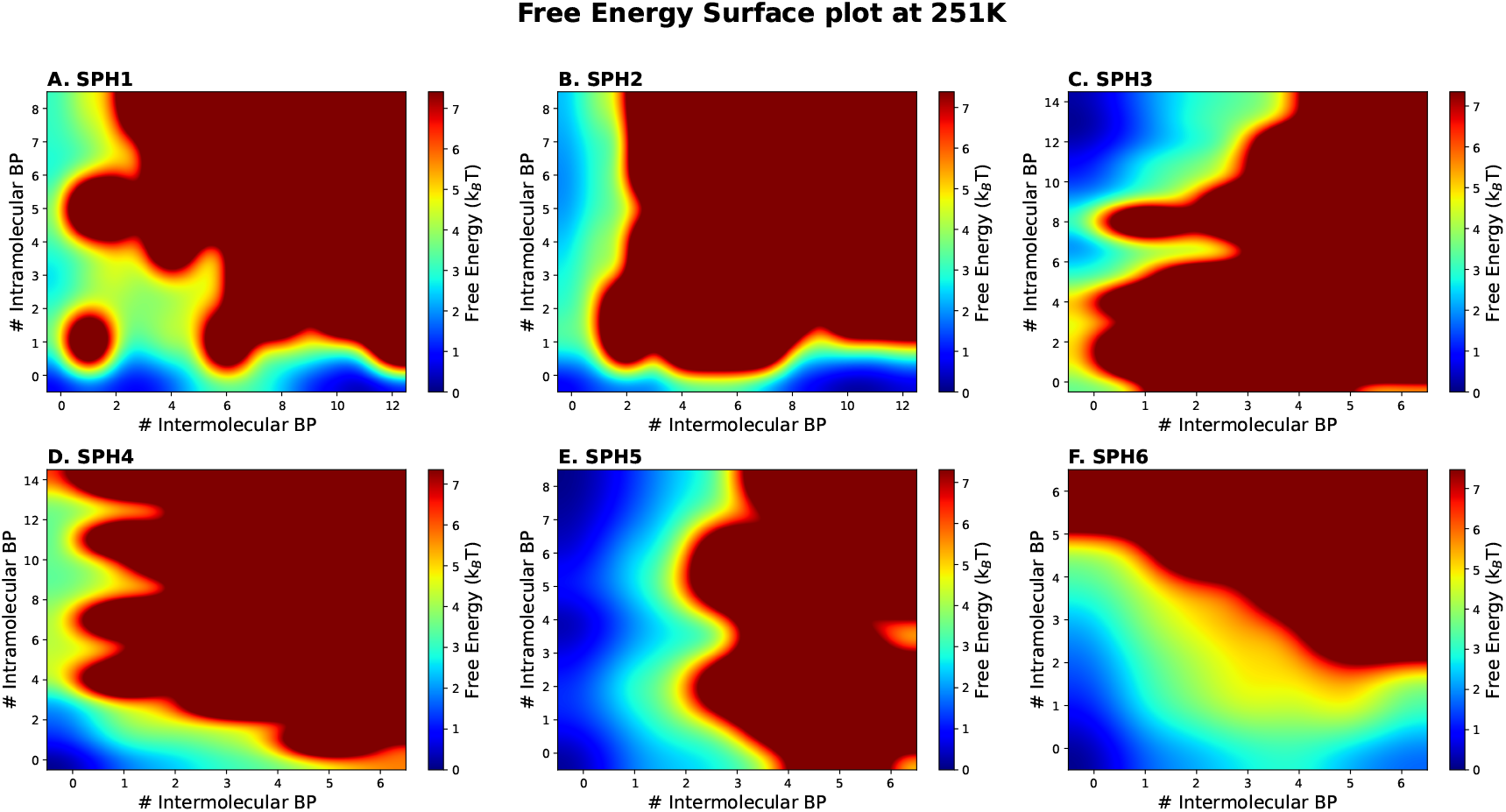
Two-dimensional free-energy surfaces *F* (*N*_intra_, *N*_inter_) for SPH1–6 at 251.1 K, computed via Boltzmann inversion of the joint probability distribution. Each panel is independently normalized so that its global minimum is zero; absolute free-energy values should therefore not be compared quantitatively across panels.

For SPH1 and SPH2 (class 1), we observe primarily competition between inter- and intra-chain base pairing, having minima at high values of one or the other. We see the global minimum at high number of intermolecular base pairs, consistent with the longer intermolecular palindrome duplex (up to 12 base pairs). Intriguingly, we find some density of points having both intra- and inter-chain base pairing for SPH1, which provides a potential pathway for binding-induced unraveling of the stem-loop structure, reminiscent of previous findings showing zipper-like unraveling of long hairpins[26].

For SPH3 (class 2), intra-chain base pairing remains high in the two-chain context and closely tracks the single-chain folding curve, while inter-chain base pairs are comparably rare. We note that inter-chain base pairing is relatively unfavorable across the full simulated temperature range (Fig. S5), likely since the dimer complex melting temperature at the simulated concentration is close to, or lower than the hairpin melting temperature. In contrast, SPH4, which has a less favorable single-chain secondary structure shows greater intermolecular base pairing, and less preference for intra-chain base paring (Figure 10D, S6).

For SPH5, the FES (Figure 10E) displays most favorable states having few intermolecular base pairs, and significant variation in the degree of intra-chain base pairing. There is also significant probability of having simultaneous intra-chain and inter-chain base pairs, owing to the design of the sequence to have the palindromic region outside of the single-chain stem-loop secondary structure. For SPH6, the FES (Figure 10F) shows few intra-chain base pairs, consistent with previous findings of low folding probability. There is significant base-pairing between chains, however, highlighting possibly the best case for forming stable inter-chain contacts, by having the palindromic regions outside any stable secondary structure.

Taken together, these results indicate that the structural context of the palindrome shapes the free-energy topology in the two-chain system, with each of the three classes producing a qualitatively distinct pattern of folding and binding accessibility in our coarse-grained model. We emphasize that these conclusions apply within our coarse-grained model and the limited set of designed test sequences; extending them to full-length transcripts is left to future work.

## Conclusions

In this work, we considered the capacity of a palindromic sequence to contribute to RNA self-association can be related to its local secondary-structure context. We refined this idea by combining a transcriptomics analysis with coarse-grained MD simulations. We found that across ∼8,000 RNA chains in the PDB and ∼533,000 transcripts in GENCODE, palindromic motifs are common. Indeed, the human transcriptome shows an enrichment of long palindromes (≳ 20 nt) relative to a random expectation. Palindromic content is broadly similar across RNA biotypes. In structurally resolved RNAs, however, most palindromic residues are sequestered within intramolecular base pairs , with fewer residues fully nested in stem-loop structures or in unpaired non-loop regions.

To examine the thermodynamic implications of this distribution, we designed six short test sequences (SPH1–6) that place a single extended palindromic region into each of the three structural classes. We tested these sequences using T-REMD simulations of both monomers and dimers. The signs of the dimer-model coefficients are consistent with the picture in which existing intramolecular structure penalizes intermolecular binding. The two-dimensional free-energy surfaces further highlight this pattern, with each class producing a qualitatively distinct landscape of folding and binding competition. Taken together, these results are consistent with the working hypothesis that the structural context of a palindrome, and not its sequence alone, is informative about the propensity of the parent RNA to self-associate in our model system.

Importantly, the insights from this work are easily implemented into high-throughput screening and transcriptomics studies. They can further be leveraged to build more sophisticated models for self-interaction preference of RNA sequences. We anticipate this will aid in further development of RNA-based therapeutics, and identification of RNA-based targets toward biomedical applications.

## Author contributions

Changhwan Ahn: Data curation (equal), Formal analysis (equal), Investigation (equal), Methodology (equal), Software (lead), Validation (equal), Visualization (lead), Writing – original draft (lead), Writing – review & editing (equal). Haebin Park: Data curation (supporting), Formal analysis (supporting), Investigation (supporting), Visualization (supporting), Writing – original draft (supporting), Writing – review & editing (supporting). Vailankanni Rodrigues: Conceptualization (lead), Data curation (equal), Investigation (supporting), Validation (supporting), Writing – review & editing (supporting). Gregory L. Dignon: Conceptualization (lead), Data curation (supporting), Formal analysis (supporting), Funding acquisition (lead), Investigation (equal), Methodology (equal), Project administration (lead), Resources (lead), Software (supporting), Supervision (lead), Validation (equal), Visualization (supporting), Writing – review & editing (lead).

## Acknowledgments

The authors would like to thank Wilma Olson, Jerelle Joseph, Patrick McCall, and Ben Schuster for helpful conversations.

## Data and code availability

Scripts used for palindrome detection and structural classification, together with data from figures, and list of pdb ids are available at https://github.com/Dignon-Lab/RNA_Palindromes_Paper_Figures. The GENCODE v49 human transcriptome is publicly available from https://www.gencodegenes.org/human/release_49.html; the PDB structures used in this work are publicly available from https://www.rcsb.org/.

