## Supporting_Information for "Palindrome Presence, Location, and Structural Context Are Predictive of RNA Self-Association"

The following supplementary figures provide additional statistical analyses supporting the bioinformatic analyses of the main text, and complete temperature-resolved free energy surface (FES) data supporting the molecular dynamics results of the main text. Figure SI 1 presents the quantitative statistical validation of palindrome distributions across PDB structural subtypes. Figure SI 2 illustrates the coarse-grained bead model representation of the SARS-CoV-2 RNA fragment used for structural classification visualization. Figures SI 3–SI 8 present the complete temperature-dependent FES panels for all six SPH sequences across all 16 T-REMD replicas, from which the reference temperature landscapes in Fig. **??** were extracted.

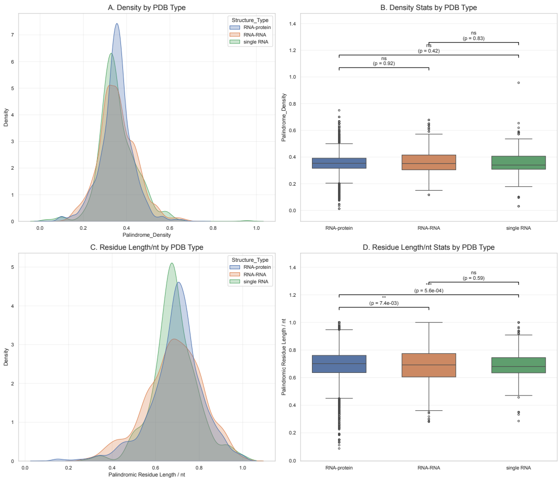


Figure SI 1: **Quantitative statistical comparison of palindrome metrics across PDB structural subtypes.** (A) Probability density distributions of palindrome count density (count/nt) for RNA–protein complexes ($N$ = 6,440), RNA–RNA complexes ($N$ = 1,137), and single RNA structures ($N$ = 485). (B) Box plots of palindrome density across PDB subtypes. Statistical significance was assessed using two-sided Mann-Whitney U tests; pairwise $p$-values are annotated above each comparison bracket (ns: not significant). No statistically significant difference in palindrome count density is observed across interaction modes, indicating that the total number of palindromic motifs per nucleotide is independent of the structural context. (C) Probability density distributions of palindromic residue length per nucleotide (fraction) across PDB subtypes. (D) Box plots of palindromic residue length fraction across PDB subtypes. In contrast to density, the palindromic residue fraction differs significantly between RNA–protein and RNA–RNA complexes ($p = 7.4 \times 10^{-3}$) and between RNA–protein and single RNA structures ($p = 5.6 \times 10^{-4}$), while RNA–RNA complexes and single RNA structures show no significant difference ($p = 0.59$). This pattern indicates that RNA molecules engaged in RNA–RNA interactions invest a larger fraction of their sequence length in palindromic motifs than those engaged exclusively with proteins, consistent with the structural availability hypothesis.

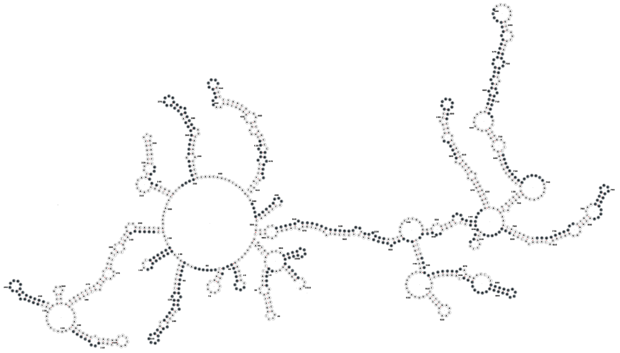

Figure SI 2: **Coarse-grained bead representation of the SARS-CoV-2 genomic RNA fragment used for structural classification.** The first 1,000 nucleotides of the SARS-CoV-2 genome are shown in the one-bead-per-nucleotide coarse-grained representation, in which each bead corresponds to a single phosphate-centered interaction site. The secondary structure connectivity is taken from the structure predicted with the RNAfold web server for the same sequence shown in Fig. **??**. This representation illustrates the spatial arrangement of the RNA backbone and the relative positions of the palindromic regions identified in the bioinformatic analysis. The overall architecture highlights the coexistence of compact hairpin domains (Class 1), long-range stem-pairing regions (Class 2), and flexible single-stranded segments (Class 3) within a single continuous RNA molecule, validating the structural diversity captured by our three-class classification scheme.

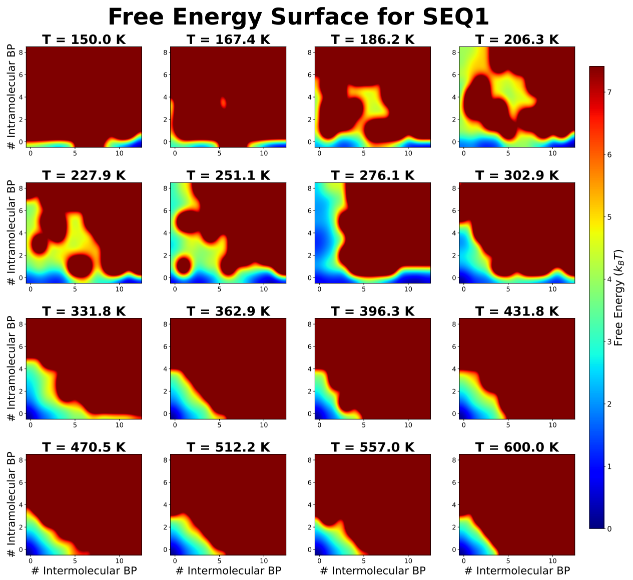


Figure SI 3: **Temperature-resolved free energy surfaces for SPH 1 across all 16 T-REMD replicas.** Two-dimensional free energy landscapes $F(N_{\text{intra}}, N_{\text{inter}})$ computed via Boltzmann inversion, $F = -k_B T \ln P$, for SPH 1 at all 16 replica temperatures spanning 150.0 K to 600.0 K. The $x$-axis and $y$-axis represent the number of intermolecular and intramolecular base pairs, respectively. Blue regions ($F \approx 0\ k_B T$) indicate thermodynamically stable states; dark red regions ($F \geq 7\ k_B T$) indicate inaccessible states. At low temperatures (150–227 K), the global free energy minimum is sharply localized along the intermolecular axis ($N_{\text{inter}} \approx$ 8–12, $N_{\text{intra}} \approx 0$), confirming that the fully bound dimer state is strongly preferred. As temperature increases (251–330 K), the dimer basin broadens and the landscape transitions through an arc-shaped low-energy pathway. Above 360 K, the system progressively accesses the fully unfolded, unbound state at the origin. The intramolecular folding basin (high $N_{\text{intra}}$, low $N_{\text{inter}}$) remains thermodynamically inaccessible at all temperatures, consistent with the near-complete suppression of intramolecular folding in each chain of the two-chain simulation observed in Fig. **??**A. The reference panel at 251.1 K corresponds to Panel A in Fig. **??**.

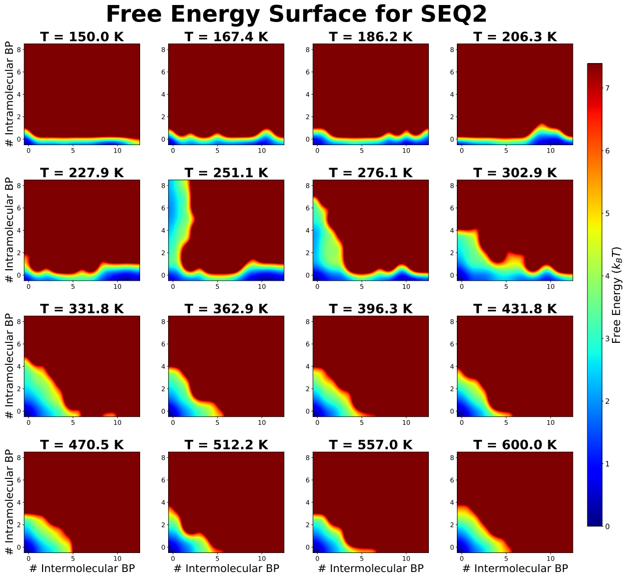


Figure SI 4: **Temperature-resolved free energy surfaces for SPH 2 across all 16 T-REMD replicas.** Two-dimensional free energy landscapes $F(N_{\text{intra}}, N_{\text{inter}})$ for SPH 2, displayed using the same format and color scale as Fig. SI 3. SPH 2 exhibits qualitatively similar behavior to SPH 1, with the global free energy minimum localized along the intermolecular axis at low temperatures. However, the dimer basin is narrower and shallower compared to SPH 1 at equivalent temperatures (e.g., 251.1 K), consistent with the modestly reduced intermolecular contact fraction observed in Fig. **??**B. This difference reflects the asymmetric tail placement of SPH 2, in which the stem is positioned closer to one terminus of the sequence, reducing the effective palindrome contact area available for intermolecular association. The intramolecular folding basin remains inaccessible across all temperatures, confirming complete binding dominance in the two-chain context. The reference panel at 251.1 K corresponds to Panel B in Fig. **??**.

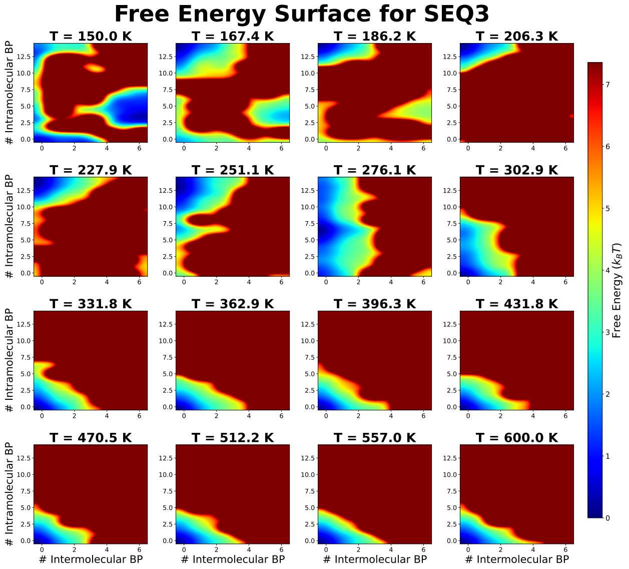


Figure SI 5: **Temperature-resolved free energy surfaces for SPH 3 across all 16 T-REMD replicas.** Two-dimensional free energy landscapes $F(N_{\text{intra}}, N_{\text{inter}})$ for SPH 3, displayed using the same format and color scale as Fig. SI 3. SPH 3 exhibits a qualitatively distinct landscape from the Hairpin-class sequences. At low temperatures (150–206 K), a characteristic vertical ladder of discrete free energy minima is visible along the $N_{\text{intra}}$ axis at $N_{\text{inter}} \approx 0$, corresponding to a series of partially folded intermediate states accessible within the comb architecture. The global minimum resides at the fully folded conformation ($N_{\text{intra}} \approx$ 12–14) and the intermolecular basin remains entirely forbidden across all temperatures, confirming that the long-range comb scaffold functions as a near-impenetrable thermodynamic lock. This is consistent with the negligible intermolecular base pairing observed in Fig. **??**C and with a $T_{m,\text{dimer}}$ at or below 150.0 K, the lowest T-REMD replica temperature, indicating that $f_{\text{inter}}$ never reaches its half-maximum within the simulated temperature range. The reference panel at 251.1 K corresponds to Panel C in Fig. **??**.

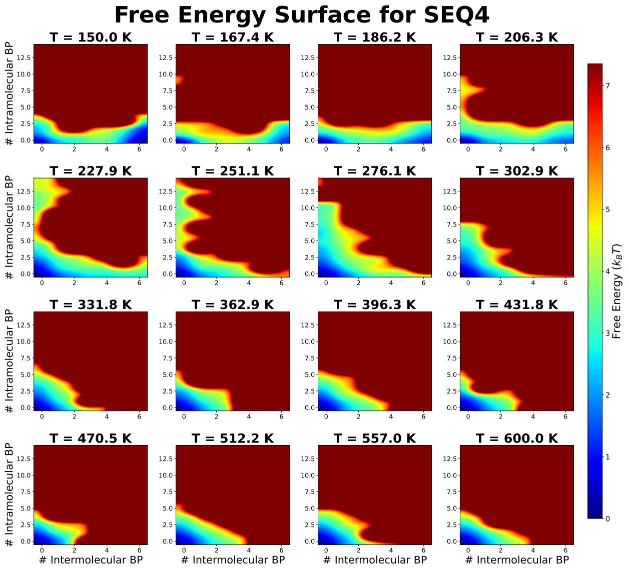


Figure SI 6: **Temperature-resolved free energy surfaces for SPH 4 across all 16 T-REMD replicas.** Two-dimensional free energy landscapes $F(N_{\mathrm{intra}}, N_{\mathrm{inter}})$ for SPH 4, displayed using the same format and color scale as Fig. SI 3. Compared to SPH 3, the introduction of a second single-nucleotide bulge in SPH 4 produces a fundamentally different landscape topology. At low temperatures (227–276 K), two distinct low-energy basins coexist: one along the intramolecular folding axis (high $N_{\mathrm{intra}}$, low $N_{\mathrm{inter}}$) and one along the intermolecular binding axis (low $N_{\mathrm{intra}}$, high $N_{\mathrm{inter}}$), separated by a shallow free energy barrier. This bifurcated landscape is the direct mechanistic signature of the additional bulge: the second mismatch does not merely weaken the fold but fundamentally reshapes the energy landscape to support two competing stable states simultaneously. At higher temperatures ($> 330$ K), both basins merge into the unfolded, unbound origin. The reference panel at 251.1 K corresponds to Panel D in Fig. **??**.

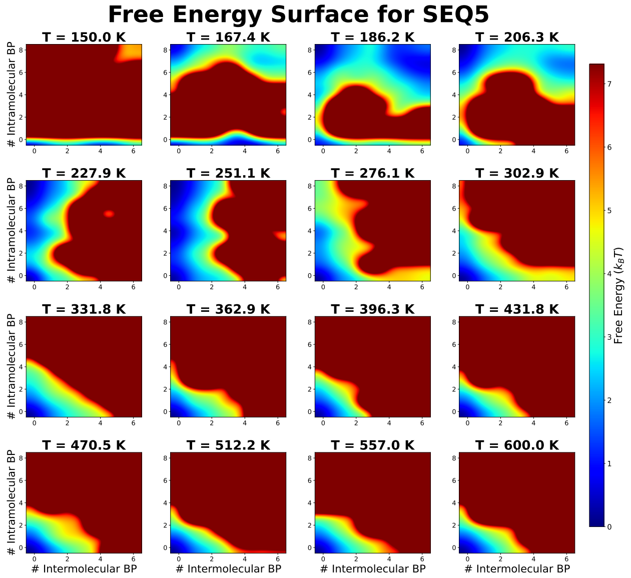


Figure SI 7: **Temperature-resolved free energy surfaces for SPH 5 across all 16 T-REMD replicas.** Two-dimensional free energy landscapes $F(N_{\text{intra}}, N_{\text{inter}})$ for SPH 5, displayed using the same format and color scale as Fig. SI 3. SPH 5 presents the most complex landscape topology of the six sequences. At low temperatures (167–251 K), a characteristic L-shaped low-energy region is visible, in which both the intramolecular folding pathway (vertical arm, $N_{\text{inter}} \approx 0$, high $N_{\text{intra}}$) and the intermolecular binding pathway (horizontal arm, $N_{\text{intra}} \approx$ 0–3, high $N_{\text{inter}}$) are simultaneously accessible and intersect at the unfolded, unbound origin. This topology directly reflects the partially exposed nature of the palindrome in SPH 5, which spans the stem-to-loop boundary: the palindromic residues can form either intramolecular or intermolecular base pairs depending on thermal fluctuations, with neither pathway strongly dominant. As the temperature increases above 302 K, the landscape collapses toward the unstructured origin. The reference panel at 251.1 K corresponds to Panel E in Fig. **??**.

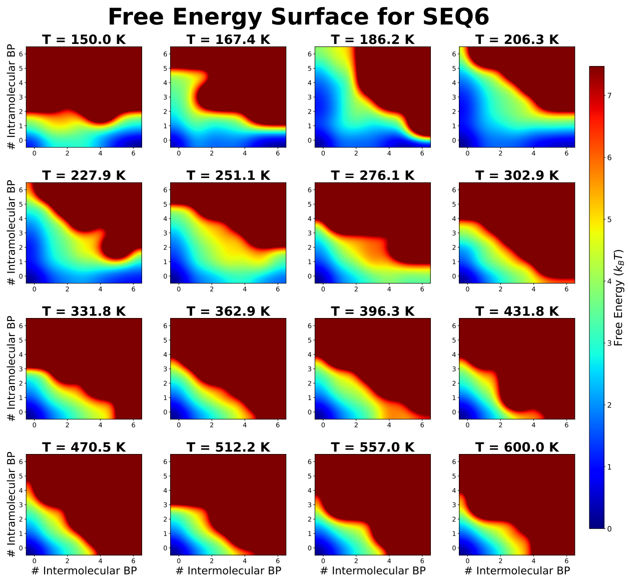


Figure SI 8: **Temperature-resolved free energy surfaces for SPH 6 across all 16 T-REMD replicas.** Two-dimensional free energy landscapes $F(N_{\text{intra}}, N_{\text{inter}})$ for SPH 6, displayed using the same format and color scale as Fig. SI 3. SPH 6 displays the most distinct landscape of all six sequences. Across the full temperature range, the landscape slopes monotonically toward high $N_{\text{inter}}$ with no discernible intramolecular folding basin at any temperature. Even at the lowest replica temperature (150.0 K), the free energy minimum resides along the intermolecular axis ($N_{\text{inter}} \approx 3$–$5$, $N_{\text{intra}} \approx 0$), with the entire intramolecular folding region ($N_{\text{intra}} > 2$) remaining inaccessible. This provides direct visual confirmation that the palindrome in SPH 6 exists in a permanently accessible state with no energetic cost to initiating intermolecular association — the weak intramolecular stability of SPH 6 found in the single-chain thermodynamic analysis ($T_{m,\text{fold}} =$ 197.8 K; Fig. **??**A). This behavior arises from the combination of a large unstructured loop ($L_{\text{loop}} =$ 15 nt) that eliminates intramolecular stability and an 8-bp palindrome that provides a substantially greater enthalpic reward for intermolecular binding than the native 4-bp intramolecular stem. The reference panel at 251.1 K corresponds to Panel F in Fig. **??**.
